# ESCRT recruitment to damaged lysosomes is dependent on the ATG8 E3-like ligases

**DOI:** 10.1101/2024.04.30.591897

**Authors:** Dale P. Corkery, Shuang Li, Deerada Wijayatunga, Laura K. Herzog, Anastasia Knyazeva, Yao-Wen Wu

## Abstract

The endosomal sorting complex required for transport (ESCRT) machinery plays an essential role in the sealing of endolysosomal membranes damaged by pathogenic, chemical or physical stress. How membrane damage is sensed by the cell and then translated into the recruitment of the ESCRT machinery is largely unknown. Here, we show that damage-dependent translocation of the autophagy ATG8 E3-like ligases to lysosomal membranes acts as the catalyst for ESCRT recruitment. Leakage of protons or calcium from perforated lysosomes induces V-ATPase-dependent or sphingomyelin-dependent recruitment of the ATG16L1-ATG5-ATG12 or TECPR1-ATG5-ATG12 E3-like complex, respectively. We show that E3-like complex-dependent recruitment of the ATG5-ATG12 conjugate to the damaged membrane is an essential prerequisite to ESCRT recruitment. At the damaged membrane ATG5-ATG12 plays both a conjugation-dependent and conjugation-independent role in stabilizing the calcium sensor, ALG-2, and recruiting the downstream repair complex. For the former scenario, we demonstrate that LC3B binds directly to ALG-2 in a Ca^2+^ dependent manner. This places the ATG8 E3-like ligases in the role of damage sensors for ESCRT-mediated membrane repair.

**GRAPHICAL ABSTRACT:** 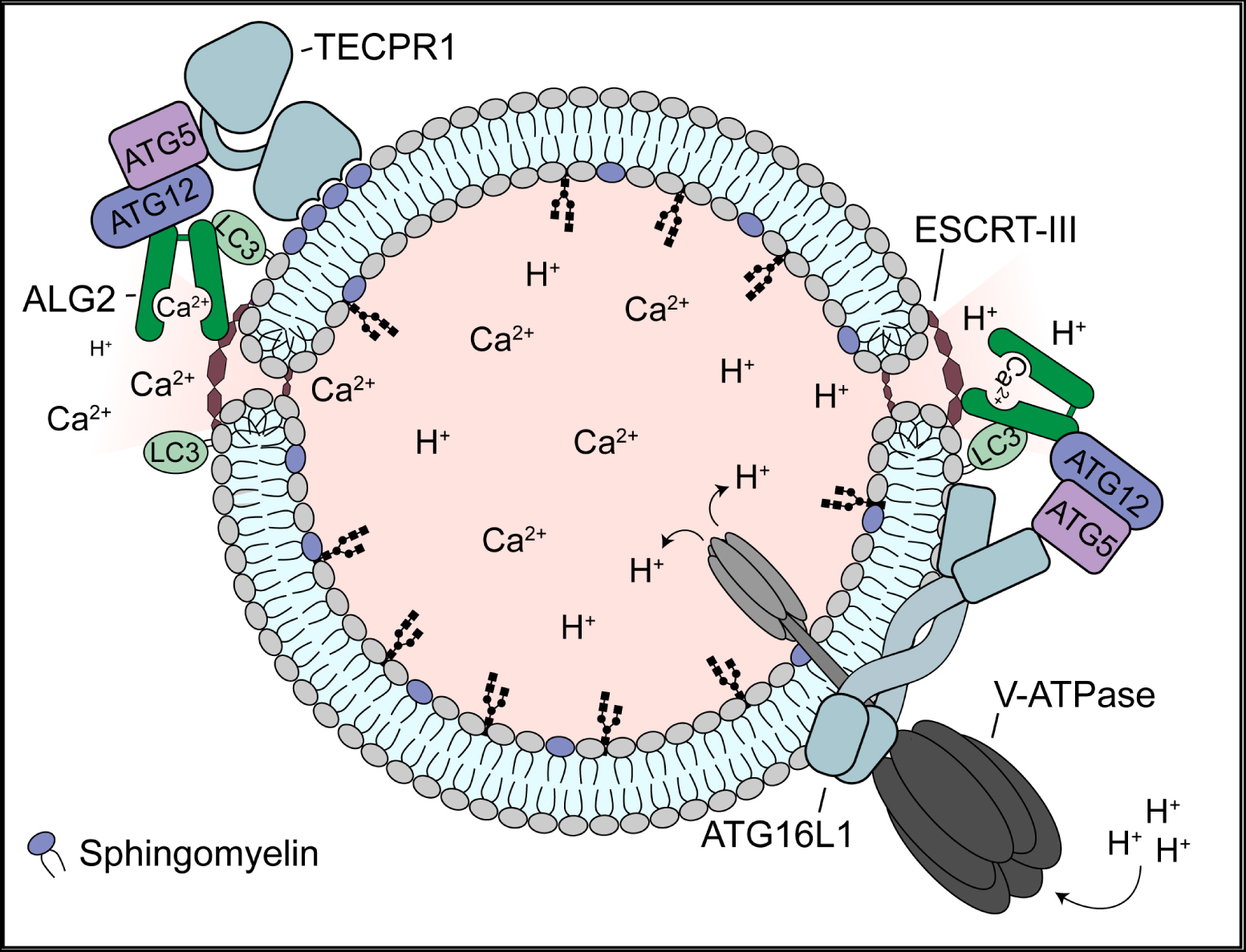

## INTRODUCTION

Membranes of the endolysosomal system face frequent damage from pathogenic, chemical or physical stress. As a result, cells have evolved sophisticated strategies to rapidly detect and repair perforated membranes. Central to this response is the endosomal sorting complex required for transport (ESCRT) machinery. This set of multisubunit protein complexes (ESCRT-0, ESCRT-I, ESCRT-II, and ESCRT-III) play an important role in membrane remodeling, and were more recently shown to play a key role in the sealing of damaged endolysosomal membranes (1, 2). The mechanism behind ESCRT-mediated membrane repair is still unknown, but is dependent on ESCRT-III filaments in combination with the ESCRT-III associated protein, ALIX, and ESCRT-I protein, TSG101 (1–3).

A fundamental question pertaining to endolysosomal membrane repair that has yet to be resolved is how the ESCRT machinery senses, and is recruited to, sites of membrane damage. Calcium efflux, in conjunction with the Ca^2+^ binding protein Apoptosis Linked Gene-2 (ALG-2), has been shown to play a central role in the sensing of damage. Lysosomal membrane damage causes Ca^2+^ to leak out of the lysosome creating a localized increase in cytosolic Ca^2+^ surrounding the damage site. Binding of ALG-2 to Ca^2+^ within this region causes ALG-2 to undergo a conformational change that promotes its interaction with ALIX (4, 5) and TSG101 (6). Failure to recruit ALG-2 to damaged lysosomal membranes prevents the subsequent recruitment of the ESCRT-III repair complex, suggesting ALG-2 is the Ca^2+^- dependent sensor that initiates ESCRT recruitment to sites of damage or osmotic stress (7). Despite ALG-2’s inherent membrane binding ability (8), recent reports have shown that ALG-2 membrane binding mutants are still recruited to lysosomes in response to damage (9) suggesting an indirect means of ALG-2 membrane recruitment. In this study, we identify the autophagy E3-like ligase complexes as an intermediate between membrane damage and ALG-2 recruitment.

Macroautophagy (hereafter autophagy) has been linked to the cellular response to membrane damage as a mechanism to sequester and degrade endomembranes that have been damaged beyond the point of repair. Extensive membrane damage will lead to endolysosomal rupture, exposing intraluminal glycans to the cytosol. The binding of glycans by a family of β-galactoside-binding lectins (galectins) serves as a platform to recruit the autophagic machinery required for sequestration of the damaged membrane into a double membraned autophagosome (10–13). A key event in autophagosome biogenesis and cargo recognition is the conjugation of autophagy-related (ATG)8 family proteins to phosphatidylethanolamine (PE) or phosphatidylserine (PS) on autophagosomal membranes (a process referred to as membrane ATG8ylation (14)). Lipidation of ATG8 proteins occurs via two ubiquitin-like ATG conjugation systems composed of core ATG genes (ATG3, ATG5, ATG7, ATG10, ATG12 and ATG16L1) (15). The ATG5-ATG12-ATG16L1 complex acts as the E3-like ligase, in which ATG16L1 recognizes target membranes and recruits the ATG5-ATG12 conjugate to catalyze the ATG8 lipidation reaction. Recently, ATG8 proteins have been shown to be conjugated to various single-membrane compartments (endolysosomal membranes, phagosomes, Golgi compartments and ER) in response to diverse stimuli. These processes are termed Conjugation of ATG8s to Single Membranes (CASM), and are characterized by the involvement of a subset of components from the autophagy machinery (16, 17). However, the functions of CASM remain largely unknown.

In the context of endolysosomal damage, ATG16L1 is recruited to damaged membranes via interaction with the Vacuolar type ATPase (V-ATPase), activated in response to damage-induced loss of the proton gradient (18, 19) (a process referred to as V-ATPase-ATG16L1- induced ATG8 lipidation (VAIL) (20)). Recently, we and others identified a second E3-like ligase complex which utilizes the autophagosome-lysosome tethering factor, Tectonin beta-propeller repeat containing 1 (TECPR1), in place of ATG16L1 (21–23). TECPR1 recognizes damaged membranes via interaction with sphingomyelin, a membrane lipid that translocates from the luminal to cytoplasmic membrane surface in response to damage (24) (a process that we call sphingomyelin-TECPR1-induced ATG8 lipidation (STIL)). Surprisingly, we found that double knockout of both E3-like ligase complexes compromised membrane repair, suggesting that the role of autophagy proteins in the cellular response to membrane damage may extend beyond autophagic removal.

In this study, we show that E3-like ligase translocation to damaged membranes is a prerequisite for ESCRT recruitment. We demonstrate that this recruitment is dependent on the ATG5-ATG12 conjugate, through both an ATG8ylation-dependent and –independent function. ATG5-ATG12 recruitment to damaged membranes, in combination with Ca^2+^, acts to stabilize ALG-2 at the damage site and promote ESCRT-mediated repair.

## RESULTS

### Loss of ATG16L1- and TECPR1-dependent ATG8 E3-like ligase complexes prevents ESCRT recruitment to damaged lysosomes

We and others recently identified a TECPR1-dependent autophagy E3-like ligase complex which functions independently of ATG16L1 to regulate unconventional ATG8 lipidation at damaged lysosomal membranes (21–23). Double knockout of both ATG16L1 and TECPR1 was shown to compromise the repair of damaged membranes, suggesting a functionally redundant role for both E3-like ligase complexes in the repair process (21). To determine the mechanism through which the E3-like ligase complexes contribute to membrane repair, we first assessed ESCRT machinery recruitment to lysosomes damaged by the lysosomal-membrane-damaging agent L-leucyl-L-leucine O-methyl ester (LLOMe), in HeLa and HEK cells deficient for ATG16L1 and/or TECPR1 (Fig 1A, B, S1A, S1B and S1C). TECPR1/ATG16L1 double knockout (E3-DKO) cells failed to recruit the ESCRT III binding protein ALIX to damaged lysosomes, despite abundant membrane damage, indicated by an accumulation of the β-galactoside-binding lectin, Galectin-3 (Gal3). Knockout of either TECPR1 or ATG16L1 alone did not prevent ALIX recruitment, consistent with a functional redundancy between the two complexes. Immunostaining for ESCRT-III complex members IST1 and CHMP2A confirmed that the ESCRT-III-dependent membrane repair complex is absent from damaged membranes in E3-DKO cells (Fig 1C and D). Importantly, the recently identified phosphoinositide-initiated membrane tethering and lipid transport (PITT) pathway for lysosomal repair (25) appears unaffected by the loss of the ATG8 E3-like ligases, as damage-induced lysosomal PI4P accumulation was evident in E3-DKO cells treated with LLOMe (Fig S2).

**Figure 1.**
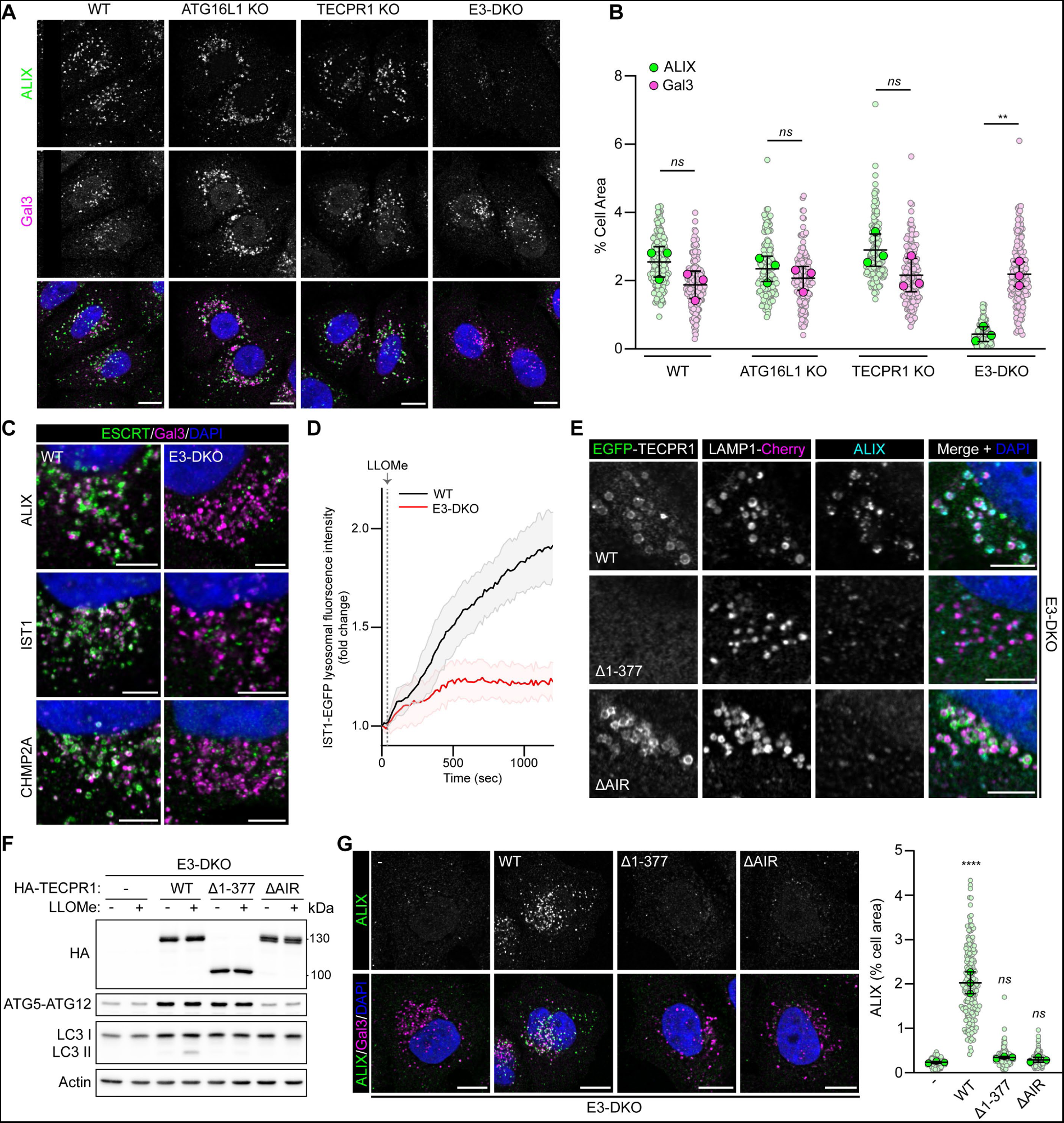
ESCRT recruitment to damaged lysosomes is impaired in cell lines deficient for the autophagy E3-like complexes. (**A**) Confocal images of HeLa WT, ATG16L1 KO, TECPR1 KO and TECPR1/ATG16L1 DKO cells treated with 1 mM LLOMe for 30 minutes. Nuclei were stained with DAPI. Scale bars = 10 µm. (**B**) Quantification of ALIX and Gal3 area from (A). Small points represent individual cells from three independent experiments. Large points represent the means of individual experiments (n = 50 cells per experiment). Bars represent the mean ± SD from the three experiments. Significance was determined from biological replicates using a one-way ANOVA with Tukey’s multiple comparisons tests. *ns* = not significant, ** p = 0.0011. (**C**) Confocal images of HeLa WT and TECPR1/ATG16L1 DKO cells treated with 1 mM LLOMe for 30 min. Scale bars = 5 µm. (D) Quantification of IST1 lysosomal recruitment in HeLa WT and E3-DKO cells co-transfected with IST1-EGFP and LAMP1-mCherry and treated with 1 mM LLOMe. Images were acquired every 15 seconds. Data are presented as mean ± SD from five independent experiments (n > 25 cells). (**E**) Confocal images of HeLa TECPR1/ATG16L1 DKO cells transfected with the indicated TECPR1 mutant and treated with 1 mM LLOMe for 30 minutes. Scale bars = 5 µm. (**F**) Western blot analysis of HeLa TECPR1/ATG16L1 DKO cells stably expressing the indicated TECPR1 mutant and treated +/- LLOMe. (**G**) Confocal images of cell lines from (E) treated with 1 mM LLOMe for 30 minutes. Quantification of ALIX area is shown to the right. Small points represent individual cells from three independent experiments. Large points represent the means of individual experiments (n = 60 cells per experiment). Bars represent the mean ± SD from the three experiments. Significance was determined from biological replicates using a one-way ANOVA with Tukey’s multiple comparisons tests. *ns* = not significant, **** p < 0.0001.

To further characterize the mechanism behind E3-like ligase-dependent ESCRT recruitment, E3-DKO addback cell lines were generated which stably express wild type (WT) TECPR1, or one of two TECPR1 mutants. TECPR1^Δ1-377^ is lacking the N-terminal dysferlin domain required for lysosomal translocation in response to membrane damage (21). TECPR1^ΔAIR^ is lacking the ATG5 interaction region (AIR) (26), preventing co-recruitment of ATG5 to the damaged membrane (Fig S3). Damage-induced lysosomal ALIX recruitment was restored with the addback of wild type TECPR1, but not with either TECPR1 mutant (Fig 1E-G), suggesting that TECPR1-dependent recruitment of ATG5 to the damaged membrane is required for subsequent ESCRT recruitment. A recent study reported that loss of ATG5 impaired ESCRT recruitment to damaged lysosomes due to an increase in the ATG12-ATG3 sidestep conjugate with an affinity for ALIX (27). The authors propose that, in the absence of its preferred conjugation partner (ATG5), ATG12 is free to form an alternative conjugate with ATG3. While we do observe reduced ATG5-ATG12 conjugate expression in E3-DKO cells (Fig S1D), the addback of TECPR1^WT^ or the lysosome-binding-deficient TECPR1^Δ1-377^ was sufficient to restore conjugate expression (Fig 1F). TECPR1^ΔAIR^ did not restore conjugate expression suggesting conjugate stability/regulation is tied to E3-like ligase complex assembly (Fig 1F). Importantly, despite its ability to restore ATG5-ATG12 conjugate expression, TECPR1^Δ1-377^ did not restore ALIX recruitment, indicating that ESCRT recruitment is dependent on ATG5 translocation to the damaged membrane.

### ESCRT recruitment to damaged lysosomes can occur independent of ATG8 lipidation

Within the trimeric E3-like ligase complexes, the ATG5-ATG12 conjugate possesses the E3- like ligase activity required for the ATG8-PE or -PS conjugation reaction (28). Our observation that ATG5 recruitment to damaged membranes is an essential prerequisite to ESCRT recruitment therefore suggests that ATG8ylation of the damaged membrane may be a contributing factor in this recruitment. To confirm this hypothesis, we evaluated the competency of ESCRT recruitment in cell lines deficient for either ATG5 or the E1-like enzyme required for the ATG5-ATG12 conjugation reaction, ATG7 (Fig 2A and B). Both cell lines are deficient for the ATG5-ATG12 conjugate (Fig 2B) and both failed to recruit ALIX to lysosomes damaged by LLOMe (Fig 2C and D). To determine if the E3-like ligase activity of the ATG5-ATG12 conjugate was required, we employed hexa-KO HeLa cells lacking the six human ATG8 paralogues (LC3A, LC3B, LC3C, GABARAP, GABARAPL1 and GABARAPL2) (ATG8KO) (29), or HeLa cells deficient for the cysteine proteases required for ATG8 processing, ATG4 A/B/C/D (30). Both cell lines continue to express the ATG5-ATG12 conjugate (Fig 2B) but are unable to generate lipidated ATG8. Surprisingly, we observed significant lysosomal ALIX recruitment in both cell lines following treatment with LLOMe (Fig 2C and D). These data suggest that, despite the requirement for ATG5-ATG12 conjugation, ATG5’s role in ESCRT recruitment is not solely linked to membrane ATG8ylation. To explore further, we stably introduced WT ATG5 or the conjugation-deficient ATG5^K130R^ mutant into ATG5KO cells (Fig 2E). Damage-induced lysosomal ALIX recruitment was restored with the addback of WT ATG5, but not with the conjugation-deficient mutant (Fig 2F), confirming a specific requirement for the ATG5-ATG12 conjugate at the damaged membrane.

**Figure 2.**
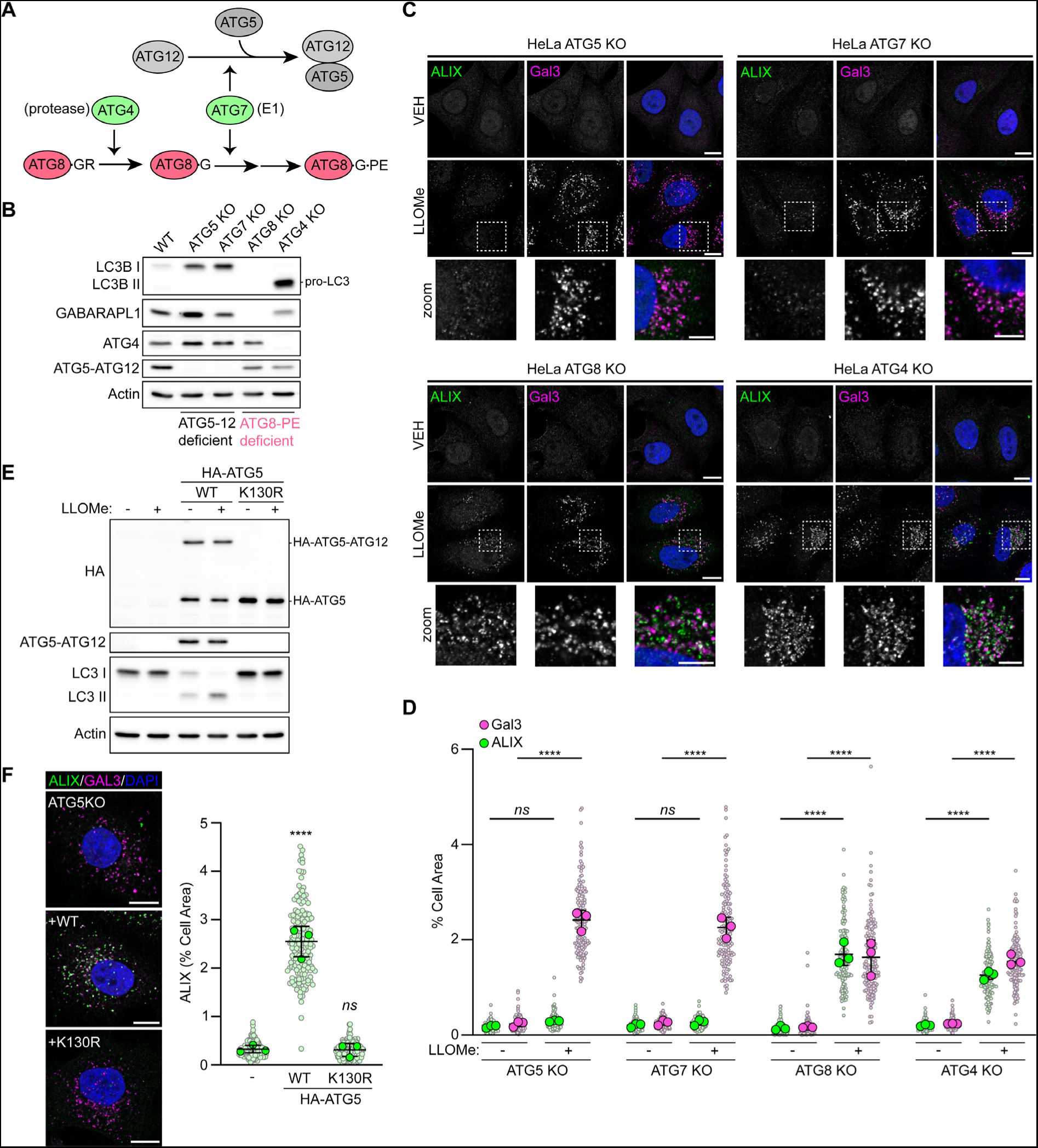
ESCRT recruitment to damaged lysosomes can occur without ATG8 lipidation. (**A**) Simplified schematic of ATG8ylation pathway. (**B**) Western blot analysis of ATG KO cell lines. (**C**) Confocal images of cell lines from (B) treated with or without 1 mM LLOMe for 30 minutes and immunostained for ALIX and Gal3. Scale bars = 10 µm for whole image and 5 µm for insets. (**D**) Quantification of ALIX and Gal3 area from (C). Small points represent individual cells from three independent experiments. Large points represent the means of individual experiments (n = 50 cells per experiment). Bars represent the mean ± SD from the three experiments. Significance was determined from biological replicates using a one-way ANOVA with Tukey’s multiple comparisons tests. *ns* = not significant, **** p = < 0.0001. (**E**) Western blot analysis of ATG5 KO cells stably expressing WT HA-ATG5 or HA-ATG5^K130R^. (**F**) Confocal images of cell lines from (E) treated with 1 mM LLOMe for 30 minutes and immunostained for ALIX and Gal3. Scale bars = 10 µm. Quantification of ALIX area is shown to the right. Small points represent individual cells from three independent experiments. Large points represent the means of individual experiments (n = 60 cells per experiment). Bars represent the mean ± SD from the three experiments. Significance was determined from biological replicates using a one-way ANOVA with Tukey’s multiple comparisons tests. *ns* = not significant, **** p < 0.0001. Comparisons against WT are shown.

### Membrane ATG8ylation is required for ESCRT-mediated repair

A recent report provided evidence suggesting that non-canonical lipidation of GABARAPs is essential for ESCRT recruitment to damaged lysosomes (31). The authors observed reduced ESCRT recruitment in GABARAP TKO cells, which they attributed to a direct interaction between GABARAPL2 and ALIX (31). In contrast to ATG5-ATG12 deficient cell lines, in which ALIX recruitment was completely abolished, we observed significant ALIX translocation to damaged lysosomes in ATG8KO cells (Fig 2C and D). To determine how this translocation compared to wild type cells, we performed immunostaining for ESCRT-III proteins IST1 and CHMP2A in HeLa WT, ATG5KO and ATG8KO cells in which lysosomes were damaged by LLOMe treatment (Fig 3A and B). Similar to ALIX, we observed increased IST1/CHMP2A puncta formation in ATG8KO cells, as compared to ATG5KO cells. However, in agreement with the above mentioned report, ESCRT translocation in ATG8KO cells was significantly less than in wild type cells. This suggests that ATG5-ATG12 plays both a conjugation-dependent and conjugation–independent role in regulating ESCRT recruitment to damaged membranes.

**Figure 3.**
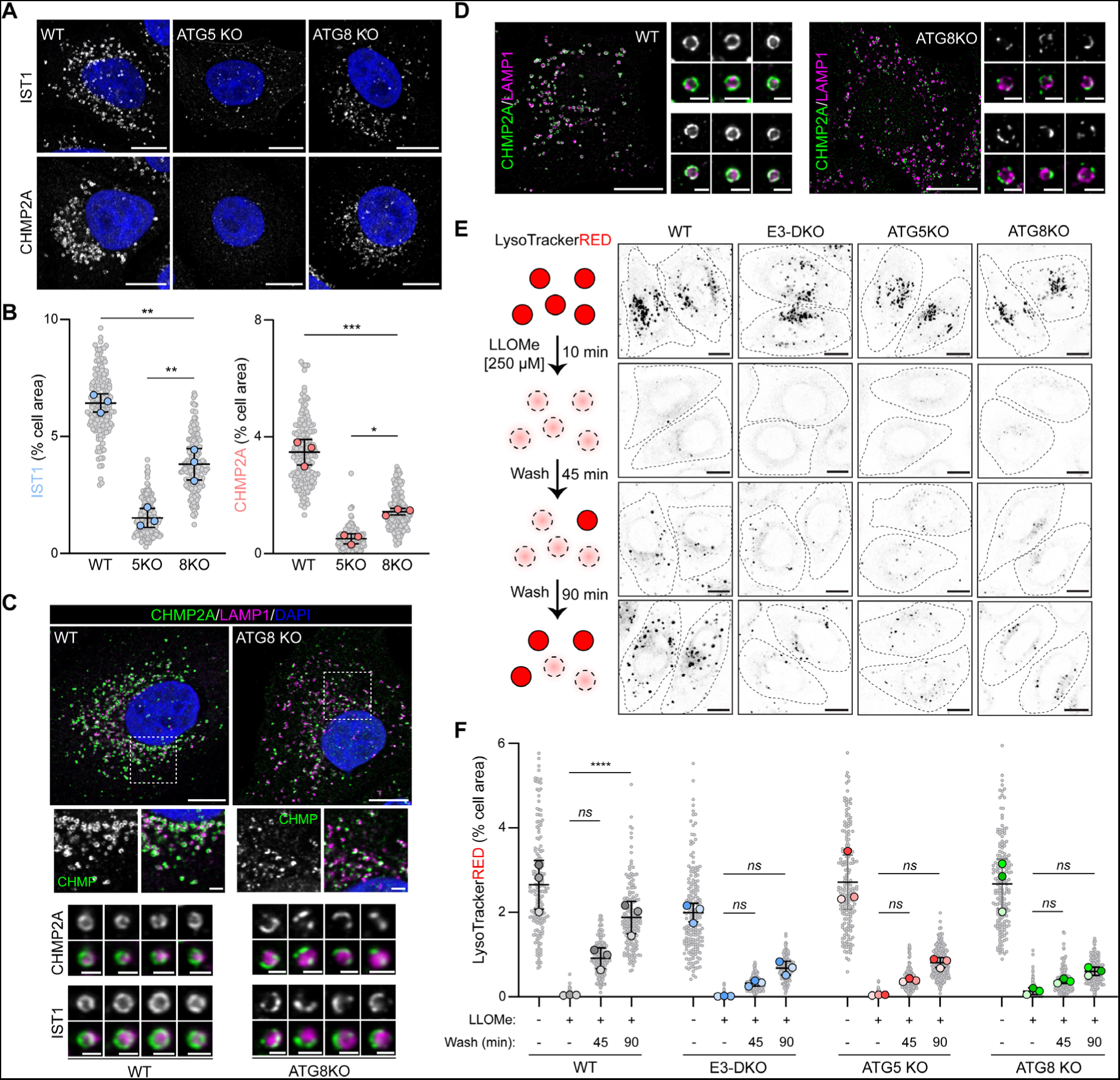
ATG8 lipidation is required for complete ESCRT recruitment and membrane repair. (**A**) Confocal images of HeLa WT, ATG5KO and ATG8KO cells treated with 1 mM LLOMe for 20 minutes. Scale bars = 10 µm. (**B**) Quantification of IST1 and CHMP2A cell area from (A). Small points represent individual cells from three independent experiments. Large points represent the means of individual experiments (n > 50 cells per experiment). Bars represent the mean ± SD from the three experiments. Significance was determined from biological replicates using a one-way ANOVA with Tukey’s multiple comparisons tests. * p = 0.0156, ** p < 0.0035, *** p = 0.0003. (**C**) Confocal images of HeLa WT and ATG8KO cells treated with 1 mM LLOMe for 20 minutes. Scale bars = 10 µm for whole cell images, 2 µm for insets, and 1 µm for individual lysosome images. (**D**) Super-resolution structured illumination images of HeLa WT and ATG8KO cells treated with 1 mM LLOMe for 20 minutes. Scale bars = 10 µm for whole cell images and 1 µm for individual lysosome images. (**E**) Representative live-cell images of HeLa cells treated as indicated and stained with LysoTracker Red. Scale bars = 10 µm. (**F**) Quantification of LysoTracker Red puncta from (C). Grey points represent individual cells from three independent experiments. Colored points represent the means of individual experiments (n = 60 cells per experiment). Bars represent the mean ± SD from the three experiments. Significance was determined from biological replicates using a one-way ANOVA with Tukey’s multiple comparisons tests. *ns* = not significant, **** p < 0.0001.

To assess the impact of membrane ATG8ylation on ESCRT recruitment, we performed high resolution (Fig 3C) or super-resolution structured illumination (Fig 3D) imaging of the ESCRT machinery on damaged lysosomes in wild type and ATG8KO cells. In wild type cells, the ESCRT machinery appears evenly distributed on lysosomes damaged by LLOMe treatment. In contrast, cell lines deficient for ATG8ylation display a much more fragmented or incomplete ESCRT distribution, which likely explains the differences in ESCRT recruitment observed between the two cell lines (Fig 3A and B). To determine if this change in ESCRT machinery distribution affected the repair process, we performed a lysosome repair assay in E3-DKO, ATG5KO and ATG8KO HeLa cells. Lysosomes were loaded with LysoTrackerRED and pulsed with 250 µM LLOMe for 10 minutes to induce damage. LLOMe was washed away and cells were allowed to recover in LysoTrackerRED containing media for 45 or 90 minutes. Restoration of LysoTrackerRED staining was used as an indicator of successful lysosome repair (Fig 3E). As previously reported, double knock out of the autophagy E3-like ligases significantly impaired lysosome recovery after damage (Fig 3F) (21). A similar impairment was observed in ATG5KO cells, consistent with our observation that E3-like ligase-dependent ATG5-ATG12 recruitment is a prerequisite for ESCRT assembly. Interestingly, lysosomal recovery in ATG8KO cells was as inefficient as in ATG5KO or E3-DKO cells, suggesting that, despite significant ESCRT recruitment, membrane ATG8ylation plays a functional role in the repair process.

### ATG8 E3-like ligases stabilize ALG-2 at damaged membranes

ALG-2 senses Ca^2+^ released into the cytosol from perforated lysosomes and recruits the ESCRT machinery to sites of damage via direct interaction with ALIX and TSG101 (7, 8). To determine if E3-like ligase recruitment to damaged membranes influences ALG-2, HeLa WT, ATG16L1KO, TECPR1KO, E3-DKO, ATG5KO and ATG8KO cells were treated with LLOMe and immunostained for ALG-2 (Fig 4A). Consistent with previous reports, we observed ALG-2 accumulation at damaged lysosomes in wild type cells (1). Knockout of ATG16L1 or TECPR1 alone had no impact on ALG-2 recruitment, while knockout of both E3-like ligases (or ATG5) blocked ALG-2 translocation to damaged membranes. This recruitment was shown to be dependent on the ATG5-ATG12 conjugate as addback of wild type ATG5 to ATG5KO cells restored ALG-2 translocation, while addback of the conjugation-deficient ATG5^K130R^ did not (Fig 4B). LLOMe-induced ALG-2 puncta were enriched for ATG5 (Fig 4C) further suggesting that E3-like ligase translocation to damaged membranes acts to recruit ALG-2. In the absence of ATG8s, ALG-2 recruitment to damaged lysosomes was impaired (Fig 4A), suggesting that membrane ATG8ylation contributes to ALG-2 recruitment. To determine if the recruitment was mediated by direct interaction between ALG-2 and ATG8, we performed an *in vitro* interaction assay using purified ALG-2 and LC3B (Fig 4D and E). LC3B was shown to interact with ALG-2 in a Ca^2+^-dependent manner suggesting that membrane ATG8ylation is one of likely several contributing factors to ALG-2 stabilization at the damaged membrane.

**Figure 4.**
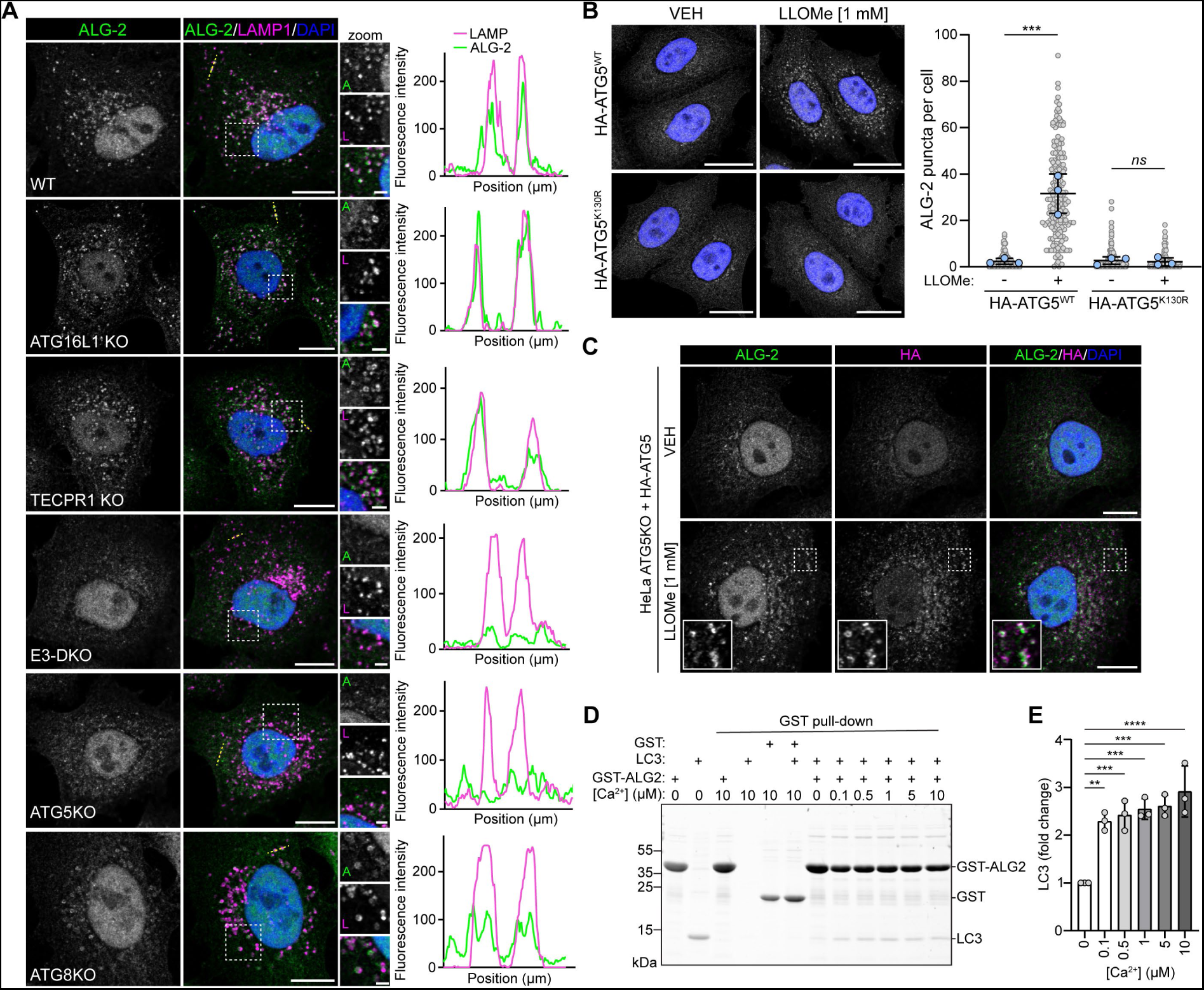
ALG-2 recruitment to damaged membranes is dependent on the E3 ligases. (**A**) Confocal images of HeLa WT, ATG16L1KO, TECPR1KO, E3-DKO, ATG5KO and ATG8KO cells treated with 1 mM LLOMe for 15 minutes. Scale bars = 10 µm for whole cell images and 2 µm for insets. Corresponding fluorescence intensity profiles are shown to the right (location marked by a yellow line on the cell image). (**B**) Confocal images of ATG5 KO cells stably expressing WT HA-ATG5 or HA-ATG5^K130R^ treated with or without LLOMe for 15 minutes. Scale bars = 20 µm. Quantification of cytosolic ALG-2 puncta is shown to the right. Small grey points represent individual cells from three independent experiments. Large blue points represent the means of individual experiments (n > 57 cells per experiment). Bars represent the mean ± SD from the three experiments. Significance was determined from biological replicates using a one-way ANOVA with Tukey’s multiple comparisons tests. ns = not significant, *** p = 0.0002. (**C**) Confocal images of ATG5 KO cells stably expressing WT HA-ATG5, treated with or without LLOMe for 15 minutes. Scale bars = 10 µm. (**D**) Representative SDS-PAGE image of GST pull-down assay. (**E**) Quantification of LC3 interaction with GST-ALG2 from (D). Data are presented as mean ± SD from the three independent experiments. Significance was determined using a one-way ANOVA with Tukey’s multiple comparisons tests. ** p = 0.0017, *** p < 0.0009, **** p < 0.0001.

## DISCUSSION

In this study, we show that the ATG8 E3-like ligases function as damage sensors, playing an essential role in ESCRT machinery recruitment to damaged lysosomal membranes. Small perforations in the lysosomal membrane will result in the leakage of both protons and Ca^2+^ into the cytosol. Collapse of the proton gradient will induce assembly of the V-ATPase proton pump on lysosomal membranes for the purpose of restoring the gradient (32). V-ATPase will subsequently recruit the ATG16L1-ATG5-ATG12 E3-like ligase complex (18) to the damaged membrane. Ca^2+^ leakage will induce the scrambling of sphingomyelin (SM) from the luminal to cytoplasmic membrane surface of the lysosomal membrane (24). SM will subsequently recruit the TECPR1-ATG5-ATG12 E3-like ligase to the damaged membrane (21–23). Here we show that ATG8 E3-like ligase translocation is an essential prerequisite to ESCRT recruitment, thereby placing the ATG8 E3-like ligases in the role of damage sensors for the ESCRT-mediated membrane repair pathway. We demonstrate that ESCRT recruitment is dependent on the ATG5-ATG12 conjugate, which plays both an ATG8 lipidation-dependent and ATG8 lipidation-independent role in regulating ESCRT recruitment to the damaged membrane. In the absence of the E3-like ligases, ALG-2 recruitment is abolished, which we attribute in part to a direct interaction between ALG2 and LC3B. We note, however, that loss of ATG8’s only partially impairs ALG-2 recruitment, suggesting that ALG-2 is further stabilized through an ATG8-indepednet interaction with the E3-like ligases.

Recent studies have offered conflicting hypotheses regarding the role of autophagy proteins in ESCRT-mediated membrane repair. Loss of ATG5 has been reported to block ESCRT recruitment to damaged lysosomes due to an increase in the ATG12-ATG3 sidestep conjugate with an affinity for ALIX (27). These conclusions were drawn, in part, due to the observation that ATG5KO, but not ATG16L1KO, impaired ALIX recruitment to lysosomes damaged with LLOMe. Our recent identification of a second E3-like ligase complex (TECPR1-ATG5-ATG12) which plays a functionally redundant role in lysosome repair (21) has allowed us to expand upon these observations by demonstrating that ATG16L1- and/or TECPR1-mediated ATG5-ATG12 recruitment to the damaged membrane is an essential step in the ESCRT repair pathway. In a second report, non-canonically lipidated ATG8s were shown to support ESCRT recruitment via a direct interaction between GABARAPL2 and ALIX (31). While we report significant damage-induced lysosomal enrichment of ALIX in the absence of ATG8s (or ATG4), a comparison against wild type cells revealed that the localization of ESCRT machinery on damaged lysosomes is distorted in the absence of membrane ATG8ylation. This supports a hypothesis whereby ATG5-ATG12, via ALG-2, recruits the ESCRT machinery to the damaged membrane. After recruitment, ESCRT localization and/or function is stabilized via direct interaction with ATG8s conjugated to the damaged membrane. This study demonstrates a fundamental function of CASM involved in ESCRT-mediated membrane repair during lysosomal damage.

## MATERIALS AND METHODS

### Cells and cell culture

HeLa WT/ATG16L1KO/ATG5KO/ATG7KO cells were a kind gift from Tomatsu Yoshimori – Osaka University, Osaka, Japan (33). HeLa WT/ATG8KO/ATG4KO cells were a kind gift from Michael Lazarou – Monash University, Melbourne, Australia (29, 30). HeLa TECPR1KO/E3-DKO cells were generated by CRISPR/Cas9-mediated knockout, as described below. HEK293 WT/ATG16L1KO were a kind gift from Anne Simonsen – University of Oslo, Oslo, Norway (34). HEK293 TECPR1KO/E3-DKO cells were described previously (21). All cells were cultured in Dulbecco’s modified Eagle medium (DMEM) (Sigma Aldrich) supplemented with 10% fetal bovine serum (FBS), 1% penicillin/streptomycin, and non-essential amino acids at 37°C with 5% CO2. Cells were routinely tested for mycoplasma contamination using the LookOut mycoplasma PCR detection kit (Sigma Aldrich).

### Generation of CRISPR KO cell lines

Oligonucleotides encoding a gRNA targeting exon 3 of TECPR1 (CACGTAGACCTGGTTGTCAC) were annealed and cloned into pSpCas9(BB)-2A-Puro (PX459) V2.0, which expresses both the Cas9 enzyme and gRNA. HeLa WT and ATG16L1KO cells were transiently transfected, selected with puromycin for 48 h, and clonal cell lines isolated by limiting dilution. TECPR1-KO cells were verified by genomic PCR amplification and sequencing of TECPR1 exon 3.

### Antibodies and reagents

Antibodies used in this study were from the following sources: ALIX (634502, IF 1:100) was purchased from BioLegend. IST1 (51002-1-AP, IF 1:200), CHMP2A (10477-1-AP, IF 1:100) and ALG-2 (12303-1-AP, IF 1:50) were purchased from Proteintech. Gal3 (87985, IF 1:400), ATG12 (2010, WB 1:1000), LC3B (#2775, WB: 1:1000), GABARAPL1 (#26632, WB: 1:1000) and ATG4B (5299, WB :1000) were purchased from Cell Signaling. Beta-actin (A2228, WB: 1:10,000) was purchased from Sigma-Aldrich. HA Tag (26183, WB 1:5000, IF 1:200) was purchased from Thermo Fisher. Alexa Fluor 488/568/647 conjugated secondary antibodies for immunofluorescence were purchased from Thermo Fisher.

Reagents used in this study were from the following sources: Leu-Leu methyl ester hydrobromide (LLOMe, L7393) from Sigma Aldrich. Lysotracker Red DND-99 (L7528) and DAPI (62248) from Thermo Fisher.

### Immunoblotting

Cells were scraped from the plate in cold lysis buffer (20 mM Tris-HCl pH8, 300 mM KCl, 10% Glycerol, 0.25% Nonidet P-40, 0.5 mM EDTA, 1 mM PMSF, 1x complete protease inhibitor (Roche)), passed through a 21G needle and cleared by centrifugation (20 min/18,213 xg/4°C). Lysates were subjected to SDS-PAGE and transferred to a 0.2 µm nitrocellulose membrane (Bio-Rad) using a Trans-Blot Turbo transfer system (Bio-Rad). Membranes were blocked in 5% skim milk (in TBST) and incubated with primary antibody diluted in 5% BSA (in TBST) overnight at 4°C. HRP-conjugated secondary antibodies were diluted in 5% skim milk (in TBST) and incubated with the membrane for 1 hour at room temperature. Protein detection was carried out using chemiluminescence (Bio-Rad) and imaged using a ChemiDoc imaging system (Bio-Rad).

### Plasmids

EGFP-TECPR1 was a kind gift from Thomas Wollert – Institute Pasteur, Paris, France (35). EGFP-TECPR1^Δ1-377^ was described previously (21). EGFP-TECPR1^ΔAIR^ was derived from EGFP-TECPR1 using PCR mutagenesis (fwd: aagaccggggcgctgcagtg, rev: catgtgtaccgaggaggacaggcc). HA-TECPR1^WT/Δ1-170/ΔAIR^ were generated by PCR amplifying TECPR1 ^WT/Δ1-170/ΔAIR^ from EGFP-TECPR1 plasmids inserting an HA tag in the forward primer. LAMP1-mCherry was described previously (21). IST1-EGFP was generated by PCR amplifying IST1 from HA-IST1 (Addgene plasmid # 131619 ; http://n2t.net/addgene:131619 ; RRID:Addgene_131619) and subcloning into the EGFP-N1 (Clontech) plasmid using NheI/SalI restriction sites. GFP-P4M-SidM was a gift from Tamas Balla (Addgene plasmid # 51469 ; http://n2t.net/addgene:51469 ; RRID:Addgene_51469) (36). mCherry-ATG5 was described previously (21). HA-ATG5 was generated by PCR amplifying ATG5 from mCherry-ATG5, adding an HA tag in the forward primer. HA-ATG5^K130R^ was derived from HA-ATG5 using PCR mutagenesis (fwd: CATTTTATGTCATGTATGAGAGAAGCTGATGCTTTAAAAC, rev: GTTTTAAAGCATCAGCTTCTCTCATACATGACATAAAATG). pSpCas9(BB)-2A-Puro (PX459) V2.0 was a gift from Feng Zhang (Addgene plasmid #62988; http://n2t.net/addgene:62988 ; RRID:Addgene_62988) (37). GST-ALG2^ΔN23^ was generated by PCR amplifying truncated ALG-2 from pDNR-PDCD6 (HsCD00000837, DNASU plasmid repository (38)) and inserting into the pGATEV-His-GST expression vector (39) using NdeI/XhoI restriction sites. LC3B (1-120, G120C) was constructed by inserting LC3B into the pMAL-6xHis-MBP-HRV 3C expression vector using KflI/HindIII restriction sites. All newly generated plasmids were verified by sanger sequencing.

### Transfection

Transfection of DNA constructs was performed using X-tremeGENE HP transfection reagent (Sigma-Aldrich) according to the manufacturer’s directions. Stable cell lines were generated via PB transposition by co-transfecting pBASE transposase with the target gene containing transposon vector at a ratio of 1:3. Cells were selected in medium containing 200 µg/ml hygromycin B for 5 days before screening for transposon integration.

### Immunofluorescence and live-cell imaging

Cells were grown on no. 1.5 glass coverslips in 6 well plates. After treatment, cells were fixed in 4% paraformaldehyde for 10 min at room temperature and permeabilized in 0.25% Triton X-100 for 5 min. Cells were blocked with 5% donkey serum for 30 min followed by a 1.5 h incubation with primary antibody at room temperature. Cells incubated with Alexa Fluor conjugated secondary antibodies for 30 min at room temperature and mounted on slides using ProLong Diamond antifade mountant (Thermo Fisher).

For live-cell imaging, cells were seeded on µ-Slide 8 well slides (Ibidi) and incubated for 24 h. Imaging was performed in DMEM without phenol red (Sigma-Aldrich) and supplemented with 20 mM HEPES.

Imaging was performed on a Leica SP8 FALCON inverted confocal system (Leica Microsystems) equipped with a HC PL APO 63x/1.40 oil immersion lens and a temperature-controlled hood maintained at 37°C and 5% CO2. DAPI was excited using a 405 nm Diode laser, and EGFP/Alexa488 and mCherry/Alexa568 fluorescence were excited using a tuned white light laser. Scanning was performed in line-by-line sequential mode.

Super-resolution Images were obtained with an Elyra 7 lattice SIM microscope (Zeiss). Images were taken at 63x with a Plan-Apochromat 63x/1.40 Oil objective using 15 phases and processed for SIM2 with the Zen Black SIM Module at default settings.

### LysoTrackerRED Repair Assay

HeLa WT/E3-DKO/ATG5KO/ATG8 KO cells were seeded in µ-Slide 8 well glass bottom slides (Ibidi) and incubated for 24 h. Lysosomes were labelled with LysoTrackerRED (Thermofisher) (0.75 µl in 10 mL media) for 30 minutes. Cells were treated with 250 µM LLOMe for 10 minutes, washed, and allowed to recover for 45 or 90 minutes in the presence of LysoTrackerRED. Cells were imaged at each time point and LysoTrackerRED area was quantified using ImageJ – FIJI distribution (NIH).

### Protein expression and purification

Proteins were expressed in *E.coli* strain BL21 (DE3) cells and grown in LB broth supplemented with 100 mg/L ampicillin. Cells were grown at 37°C on a shaker (160 rpm) until the absorbance at 600 nm (OD600) reached 0.5-0.7. Protein expression was induced with 0.4 mM Isopropyl β-d-1-thiogalactopyranoside (IPTG) treatment overnight at 23°C (160 rpm). Cells were harvested by centrifugation at 5000 rpm, 4°C for 30 min, then discard the supernatant carefully and wash the cells with PBS by centrifugation.

All purification steps were performed at 4°C unless otherwise noted. Cells were resuspended in lysis buffer (50 mM Tris, 100 mM NaCl, 5 % glycerol, pH 8.0) with freshly added 1 mM phenylmethylsulfonyl fluoride (PMSF) and 2 mM β-mercaptoethanol. Resuspended cells were lysed using a cell disruptor (Constant Systems) at 27 kpsi three times. The cell lysates were further supplemented with 1% Triton X-100 and then centrifuged at 18000 rpm for 1 h. The supernatant was filtered through a 0.2 µm membrane and then loaded onto a 5 ml HisTrap HP (Cytiva) column, which was equilibrated with buffer A (50 mM Tris, 100 mM NaCl, 5% glycerol and freshly added 2 mM β-mercaptoethanol, pH 8.0) and then protein was eluted with an isocratic of buffer B (50 mM Tris, 100 mM NaCl, 5% glycerol, 500 mM Imidazole and fresh 2 mM β-mercaptoethanol, pH 8.0). The fraction of interest was concentrated with 10 kDa Amicon Ultra centrifuge filter to ∼10 ml. Finally, the protein was purified by size-exclusion chromatography (Superdex 200 pg, HiLoad 26/60) in 50 mM Tris, 100 mM NaCl, 5% Glycerol and fresh 2 mM β-mercaptoethanol, pH 8.0. Fractions containing the peak of interest were concentrated with Amicon 10 kDa to a final concentration of ∼9.4 mg/mL. 2 mM DTT were added to proteins for storage. For LC3B (1-120), the His-MBP tag was removed by adding 0.1 eq PreScission protease and dialyzed in buffer (50 mM Tris, pH 7.5, 300 mM NaCl, 10 % (v/v) glycerol) overnight at room temperature. The cleaved protein was purified on a HisTrap HP column again and the flow through fractions were collected. After concentration with 10 kDa Amicon ultra centrifuge filter, the protein was purified by size-exclusion chromatography and concentrated to a final concentration of 1.0 mg/mL with 2 mM TECP was added for storage. For His-tagged GST, His-GST-TEV-ALG2^ΔN23^ was incubated with 1% (w/w) TEV protease and dialyzed in buffer (50 mM Tris, 100 mM NaCl, 5% glycerol, 2 mM β-mercaptoethanol, pH 8.0). His-GST was purified as described above. The final concentration of his-tagged GST was 1.8 mg/mL with 2 mM DTT added for storage.

### GST pull-down assay

Purified recombinant GST or GST-ALG2 and LC3B proteins (10 µM) were mixed together in buffer A (50 mM Tris, pH 8.0, 100 mM NaCl, 5 % Glycerol) in the presence of 1 mM EDTA or 0.1-10 µM Ca^2+^ and incubated with 25 μL glutathione (GSH) high-capacity magnetic agarose beads (Sigma, G0924) for 4 h at 4°C. After washing three times with buffer A, proteins were eluted from the beads with GSH-containing buffer (50 mM Tris, pH 8.0, 100 mM NaCl, 5 % Glycerol, 20 mM GSH) in the presence of 1 mM EDTA or 0.1-10 µM Ca^2+^, and subjected to SDS-PAGE analysis.

### Quantification and statistical analysis

Data are shown as mean ± standard deviation (SD). Statistical significance was determined by one-way ANOVA with Tukey’s multiple comparisons tests, using GraphPad Prism v.10.0.0. *p < 0.05, **p < 0.01, ***p < 0.001, ns = not significant.

## ACKNOWLEDGEMENTS

This work was supported by the European Research Council (ChemBioAP), Vetenskapsrådet (Nr. 2018-04585, Nr. 2022-02932), the Knut and Alice Wallenberg Foundation and the Göran Gustafsson Foundation for Research in Natural Sciences and Medicine to Y.W.W. We thank Irene Martinez Carrasco for assistance with the super-resolution imaging, and acknowledge the Biochemical Imaging Center at Umeå University and the National Microscopy Infrastructure, NMI (VR-RFI 2019-00217) for providing assistance in microscopy.

## AUTHOR CONTRIBUTIONS

Conceptualization and formal analysis, D.P.C and Y.W; investigation and validation, D.P.C., S.L., D.W., L.K.H and A.K; writing – original draft, D.P.C; data curation; D.P.C., S.L., L.K.H. and A.K.; visualization; D.P.C; supervision, project administration and funding acquisition, Y.W.

## Figures

**Figure S1.**
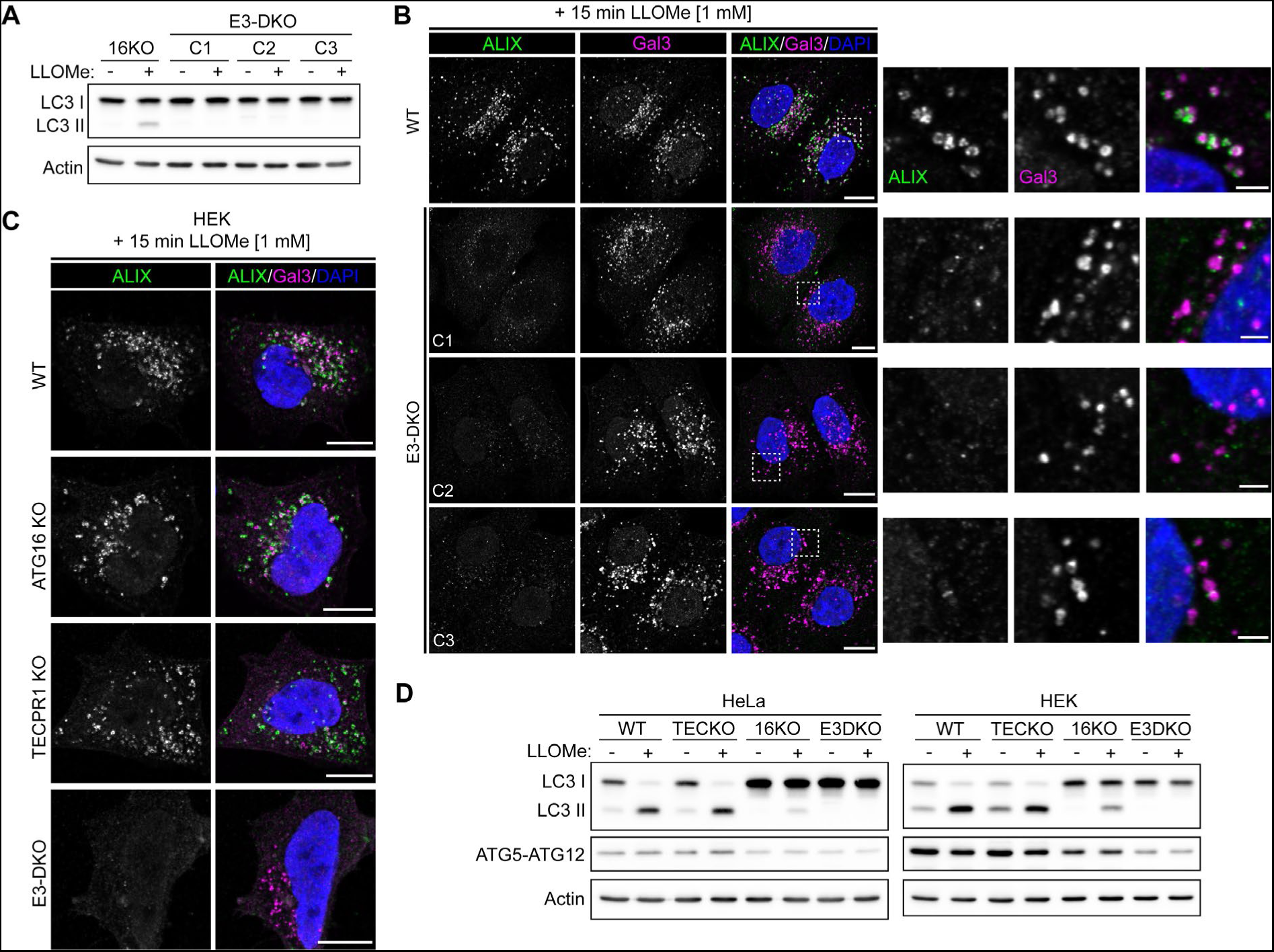
Validation of ESCRT recruitment deficiency across multiple ATG16L1/TECPR1 DKO clones and cell lines. (**A**) Western blot analysis of HeLa ATG16L1KO and three ATG16L1/TECPR1 DKO clones treated with or without 1 mM LLOMe for 30 minutes. (**B**) Confocal images of cell lines from (A) treated with 1 mM LLOMe for 15 minutes and immunostained for ALIX and Gal3. Nuclei were stained with DAPI. Scale bars = 10 µm for whole image and 2 µm for insets. (**C**) Confocal images of HEK WT, ATG16L1KO, TECPR1KO and ATG16L1/TECPR1 DKO cells treated with 1 mM LLOMe for 15 minutes and immunostained for ALIX and Gal3. Scale bars = 10 µm. (**D**) Western blot analysis of HeLa and HEK KO cell lines treated with or without 1 mM LLOMe for 30 minutes.

**Figure S2.**
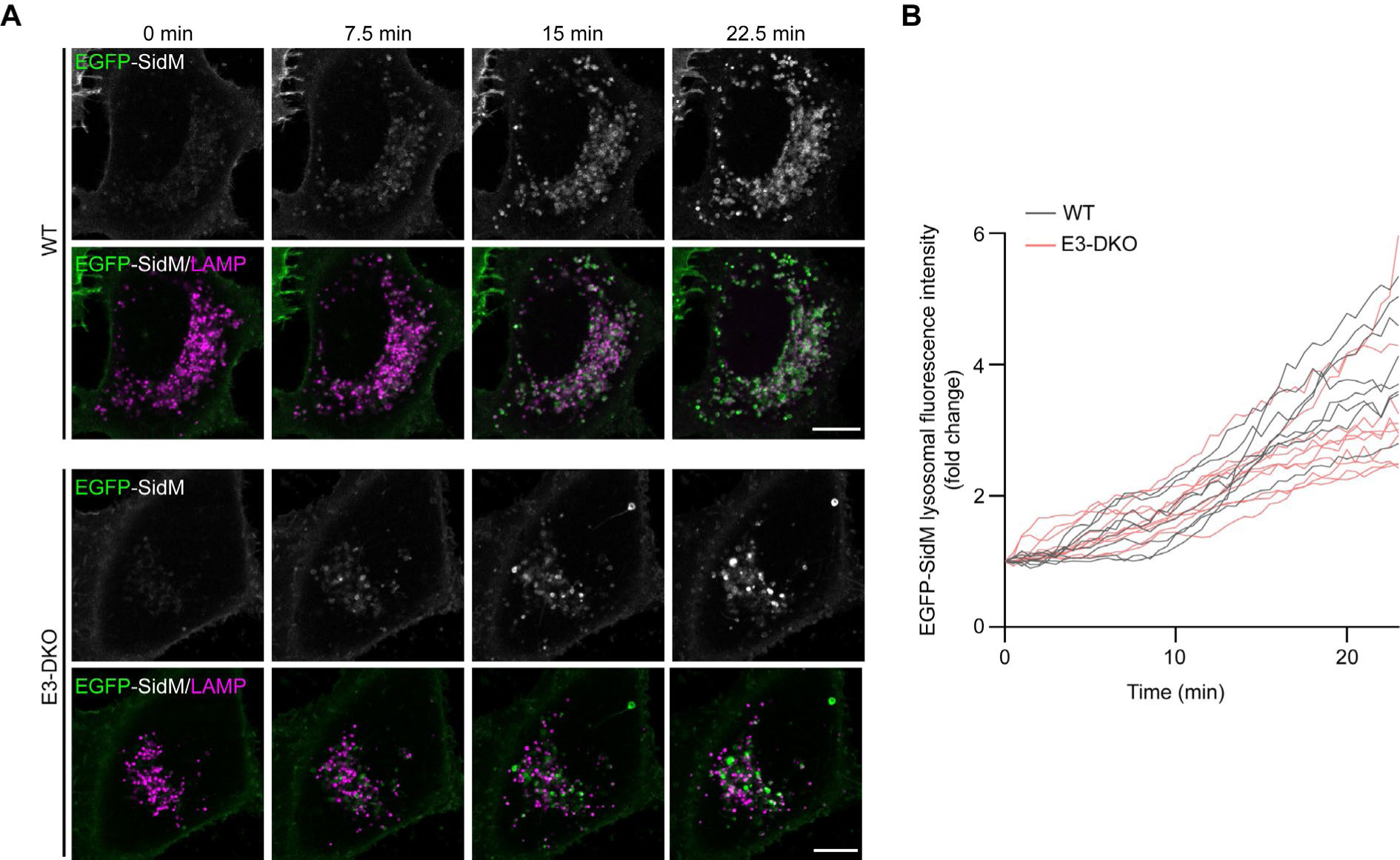
Damage-induced lysosomal PI4P accumulation is unaffected by the loss of the ATG8 E3 ligase complexes. (**A**) Representative live-cell fluorescent images of HeLa WT and ATG16L1/TECPR1 DKO cells co-transfected with EGFP-SidM (PI4P biosensor) and LAMP1-mCherry and treated with 1 mM LLOMe for the indicated time. Scale bar = 10 µM. (**B**) Quantification of the fold change in EGFP-SidM lysosomal fluorescence from (A). Lines represent individual cells from three independent experiments.

**Figure S3.**
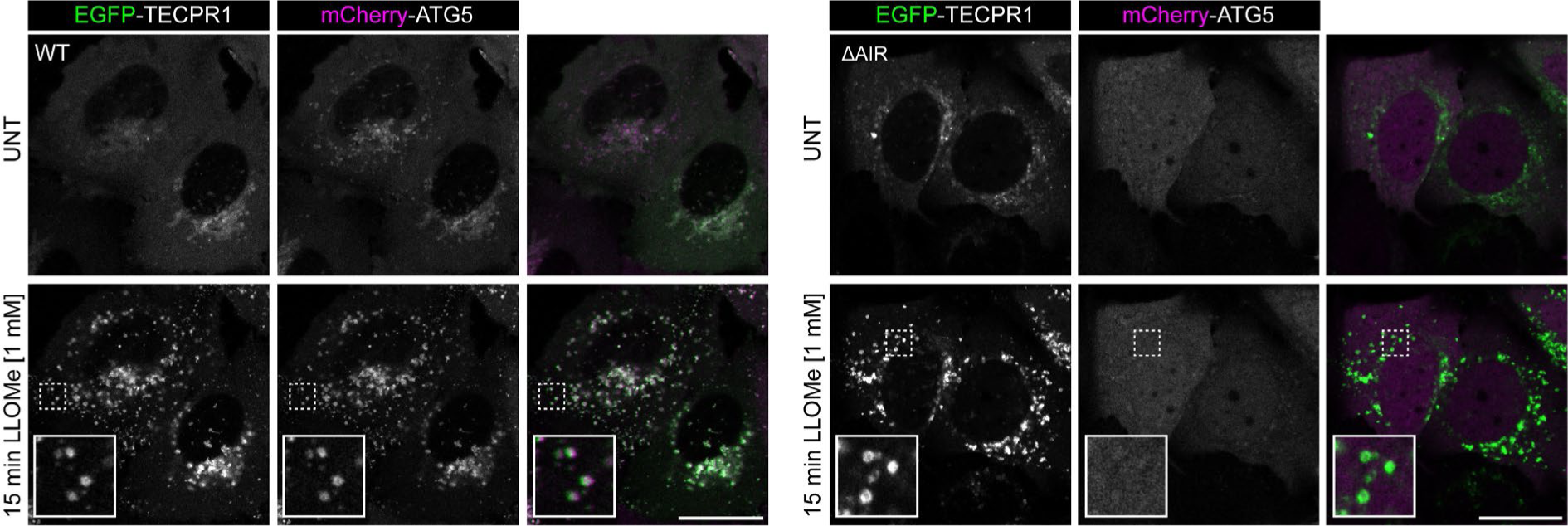
Deletion of the ATG5 interacting region of TECPR1 prevents ATG5 recruitment to lysosomes in response to damage. Representative live-cell fluorescent images of HeLa WT cells co-transfected with EGFP-TECPR1 and mCherry-ATG5 and treated with 1 mM LLOMe for 15 minutes. Scale bar = 20 µM.

## REFERENCES

1. M. L. Skowyra, P. H. Schlesinger, T. V. Naismith, P. I. Hanson, Triggered recruitment of ESCRT machinery promotes endolysosomal repair. Science 360 (2018).

2. M. Radulovic et al., ESCRT-mediated lysosome repair precedes lysophagy and promotes cell survival. EMBO J 37 (2018).

3. J. J. Chen et al., Compromised function of the ESCRT pathway promotes endolysosomal escape of tau seeds and propagation of tau aggregation. J Biol Chem 294, 18952–18966 (2019).

4. M. Missotten, A. Nichols, K. Rieger, R. Sadoul, Alix, a novel mouse protein undergoing calcium-dependent interaction with the apoptosis-linked-gene 2 (ALG-2) protein. Cell Death Differ 6, 124–129 (1999).

5. H. Suzuki et al., Structural basis for Ca2+ -dependent formation of ALG-2/Alix peptide complex: Ca2+/EF3-driven arginine switch mechanism. Structure 16, 1562–1573 (2008).

6. K. Katoh et al., The penta-EF-hand protein ALG-2 interacts directly with the ESCRT-I component TSG101, and Ca2+-dependently co-localizes to aberrant endosomes with dominant-negative AAA ATPase SKD1/Vps4B. Biochem J 391, 677–685 (2005).

7. W. Chen, M. M. Motsinger, J. Li, K. P. Bohannon, P. I. Hanson, Ca. bioRxiv (2024).

8. S. Shukla, K. P. Larsen, C. Ou, K. Rose, J. H. Hurley, In vitro reconstitution of calcium-dependent recruitment of the human ESCRT machinery in lysosomal membrane repair. Proc Natl Acad Sci U S A 119, e2205590119 (2022).

9. S. Shukla et al., Mechanism and cellular function of direct membrane binding by the ESCRT and ERES-associated Ca. Proc Natl Acad Sci U S A 121, e2318046121 (2024).

10. S. Chauhan et al., TRIMs and Galectins Globally Cooperate and TRIM16 and Galectin-3 Co-direct Autophagy in Endomembrane Damage Homeostasis. Dev Cell 39, 13–27 (2016).

11. T. L. Thurston, M. P. Wandel, N. von Muhlinen, A. Foeglein, F. Randow, Galectin 8 targets damaged vesicles for autophagy to defend cells against bacterial invasion. Nature 482, 414–418 (2012).

12. I. Paz et al., Galectin-3, a marker for vacuole lysis by invasive pathogens. Cell Microbiol 12, 530–544 (2010).

13. I. Maejima et al., Autophagy sequesters damaged lysosomes to control lysosomal biogenesis and kidney injury. EMBO J 32, 2336–2347 (2013).

14. S. Kumar, J. Jia, V. Deretic, Atg8ylation as a general membrane stress and remodeling response. Cell Stress 5, 128–142 (2021).

15. N. Mizushima, The ATG conjugation systems in autophagy. Curr Opin Cell Biol 63, 1–10 (2020).

16. L. Galluzzi, D. R. Green, Autophagy-Independent Functions of the Autophagy Machinery. Cell 177, 1682–1699 (2019).

17. J. Durgan, O. Florey, Many roads lead to CASM: Diverse stimuli of noncanonical autophagy share a unifying molecular mechanism. Sci Adv 8, eabo1274 (2022).

18. Y. Xu et al., A Bacterial Effector Reveals the V-ATPase-ATG16L1 Axis that Initiates Xenophagy. Cell 178, 552–566.e520 (2019).

19. K. Fletcher et al., The WD40 domain of ATG16L1 is required for its non-canonical role in lipidation of LC3 at single membranes. EMBO J 37 (2018).

20. T. D. Fischer, C. Wang, B. S. Padman, M. Lazarou, R. J. Youle, STING induces LC3B lipidation onto single-membrane vesicles via the V-ATPase and ATG16L1-WD40 domain. J Cell Biol 219 (2020).

21. D. P. Corkery, S. Castro-Gonzalez, A. Knyazeva, L. K. Herzog, Y. W. Wu, An ATG12-ATG5-TECPR1 E3-like complex regulates unconventional LC3 lipidation at damaged lysosomes. EMBO Rep, e56841 (2023).

22. N. Kaur et al., TECPR1 is activated by damage-induced sphingomyelin exposure to mediate noncanonical autophagy. EMBO J, e113105 (2023).

23. K. B. Boyle et al., TECPR1 conjugates LC3 to damaged endomembranes upon detection of sphingomyelin exposure. EMBO J, e113012 (2023).

24. P. Niekamp et al., Ca 2+ -activated sphingomyelin scrambling and turnover mediate ESCRT-independent lysosomal repair. Nat Commun 13, 1875 (2022).

25. J. X. Tan, T. Finkel, A phosphoinositide signalling pathway mediates rapid lysosomal repair. Nature 609, 815–821 (2022).

26. D. Chen et al., A mammalian autophagosome maturation mechanism mediated by TECPR1 and the Atg12-Atg5 conjugate. Mol Cell 45, 629–641 (2012).

27. F. Wang et al., ATG5 provides host protection acting as a switch in the atg8ylation cascade between autophagy and secretion. Dev Cell 58, 866–884.e868 (2023).

28. T. Hanada et al., The Atg12-Atg5 conjugate has a novel E3-like activity for protein lipidation in autophagy. J Biol Chem 282, 37298–37302 (2007).

29. T. N. Nguyen et al., Atg8 family LC3/GABARAP proteins are crucial for autophagosome-lysosome fusion but not autophagosome formation during PINK1/Parkin mitophagy and starvation. J Cell Biol 215, 857–874 (2016).

30. T. N. Nguyen et al., ATG4 family proteins drive phagophore growth independently of the LC3/GABARAP lipidation system. Mol Cell 81, 2013–2030.e2019 (2021).

31. M. Ogura et al., Microautophagy regulated by STK38 and GABARAPs is essential to repair lysosomes and prevent aging. EMBO Rep 24, e57300 (2023).

32. R. J. Mulligan, et al., Collapse of late endosomal pH elicits a rapid Rab7 response via V-ATPase and RILP. bioRxiv (2024).

33. S. Nakamura et al., LC3 lipidation is essential for TFEB activation during the lysosomal damage response to kidney injury. Nat Cell Biol 22, 1252–1263 (2020).

34. A. H. Lystad et al., Distinct functions of ATG16L1 isoforms in membrane binding and LC3B lipidation in autophagy-related processes. Nat Cell Biol 21, 372–383 (2019).

35. L. Wetzel et al., TECPR1 promotes aggrephagy by direct recruitment of LC3C autophagosomes to lysosomes. Nat Commun 11, 2993 (2020).

36. G. R. Hammond, M. P. Machner, T. Balla, A novel probe for phosphatidylinositol 4-phosphate reveals multiple pools beyond the Golgi. J Cell Biol 205, 113–126 (2014).

37. F. A. Ran et al., Genome engineering using the CRISPR-Cas9 system. Nat Protoc 8, 2281–2308 (2013).

38. C. Y. Seiler et al., DNASU plasmid and PSI:Biology-Materials repositories: resources to accelerate biological research. Nucleic Acids Res 42, D1253–1260 (2014).

39. A. Kalinin et al., Expression of mammalian geranylgeranyltransferase type-II in Escherichia coli and its application for in vitro prenylation of Rab proteins. Protein Expr Purif 22, 84–91 (2001).

